# Human CardioChimeras: Creation of a Novel ‘Next Generation’ Cardiac Cell

**DOI:** 10.1101/796870

**Authors:** Fareheh Firouzi, Sarmistha Sinha Choudhury, Kathleen Broughton, Adriana Salazar, Mark A Sussman

**Affiliations:** San Diego State University, Department of Biology and Integrated Regenerative Research Institute, San Diego, California

**Keywords:** cell fusion, cardiac interstitial cells, mesenchymal stromal cells, cardiac, human

## Abstract

**Background:** CardioChimeras (CCs) produced by fusion of murine c-kit^+^ cardiac interstitial cells (cCIC) with mesenchymal stem cells (MSCs) promote superior structural and functional recovery in a mouse model of myocardial infarction (MI) compared to either precursor cell alone or in combination. Creation of human CardioChimeras (hCC) represents the next step in translational development of this novel cell type, but new challenges arise when working with cCICs isolated and expanded from human heart tissue samples. The objective of the study was to establish a reliable cell fusion protocol for consistent optimized creation of hCCs and characterize fundamental hCC properties.

**Methods and Results:** Cell fusion was induced by incubating human cCICs and MSCs at a 2:1 ratio with inactivated Sendai virus. Hybrid cells were sorted into 96-well microplates for clonal expansion to derive unique cloned hCCs, which were then characterized for various cellular and molecular properties. hCCs exhibited enhanced survival relative to the parent cells and promoted cardiomyocyte survival in response to serum deprivation in vitro.

**Conclusions:** The generation of hCC is demonstrated and validated in this study, representing the next step toward implementation of a novel cell product for therapeutic development. Feasibility of creating human hybrid cells prompts consideration of multiple possibilities to create novel chimeric cells derived from cells with desirable traits to promote healing in pathologically damaged myocardium.

**Clinical Perspective:**

- “Next generation” cell therapeutics will build upon initial findings that demonstrate enhanced reparative action of combining distinct cell types for treatment of cardiomyopathic injury.
- Differential biological properties of various cell types are challenging for optimization of delivery, engraftment, persistence, and synergistic action when used in combination.
- Creation of a novel hybrid cell called a CardioChimera overcomes limitations inherent to use of multiple cell types.
- CardioChimeras exhibit unique properties relative to either parental cell anticipated to be advantageous in cellular therapeutic applications.
- CardioChimeras have now been created and characterized using cells derived from human heart tissue, advancing initial proof of concept previously demonstrated with mice.
- CardioChimeras represent an engineered solution that can be implemented as a path forward for improving the outcome of myocardial cell therapy.

## Introduction

Beneficial, albeit modest, effects of cell therapy for treatment of myocardial damage have been established in extensive preclinical studies^1,2^ as well as early clinical trial testing^3,4,5^. Improvement of efficacy remains a primary focus of the cell therapy field with ongoing studies attempting to define whether there is an optimal cell type or even a combination of cell types delivered simultaneously to promote reparative remodeling^6,7^. The concept of creating an ‘enhanced’ cell with augmented biological properties to promote healing remains a longstanding interest of our group. Simply stated, we have sought to create ‘unnatural solutions’ to the natural limitations of myocardial repair by ex vivo modification resulting in engineered cell products derived from c-kit+ cardiac interstitial cells (cCICs) possessing enhanced survival, persistence, proliferation, and many other desirable characteristics that deliver greater recovery from myocardial injury than corresponding originating parental cells^8,9,10,11^. One example of this philosophy is the CardioChimera, named for the fusion of cCICs and mesenchymal stem cells (MSCs) together into a single hybrid cell that can be expanded in culture and used for adoptive transfer therapy to mitigate myocardial infarction injury^8^. The seminal study of CardioChimeras was performed using a homotypic murine model system with delivery murine CardioChimeras and results demonstrated superiority of the CardioChimeras over either cCICs alone, MSCs alone, or the combination of cCICs and MSCs delivered together as a single cell suspension. These promising findings prompted subsequent studies to translate the murine findings into the human context but, working with myocardial-derived human cells presents a new set of challenges to be overcome.

As opposed to murine cCICs and MSCs obtained from 2-3-month-old hearts with high yield quantities^8^, human cardiac cells are derived from small ventricular biopsy specimens^12^. Limited source material results in lower yield of human myocardial-derived cells, and the relatively slow proliferation of myocardial-derived human cells relative to mouse counterparts^12,13^ further hampers therapeutic manipulations. Furthermore, replicative senescence occurs much earlier in culture passaging of human cells, whereas murine cardiac cells demonstrate significantly extended passaging capability^8,14,15^. Another technical challenge posed by inherent species-specific biology are differences between murine and human cells in response to experimental interventions^16^. Murine cells readily fuse and form polyploid hybrid cells^8^ whereas human cells are much less tolerant of polyploidy leading to genomic instability^17,18^ that necessitates rigorous optimization of culture conditions to overcome impaired expansion following fusion. Efficient and reproducible creation of CardioChimeras is facilitated by use of murine cardiac cell populations from syngeneic healthy young donors^8^ unlike the individual patient-specific nature of human myocardial derived cells with unavoidable sample characteristic variability^14^. Collectively, these challenges prompted this study to demonstrate feasibility and reproducibility of human CardioChimera (hCC) generation.

Concurrent isolation of three distinct CIC types comprised of cCIC, MSC, and endothelial progenitor cells (EPC) from a single human myocardial tissue sample with optimized culture condition was developed by our group^12^. Therefore, this protocol was employed to obtain human cCIC and MSC for subsequent fusion proof of concept using low passage neonatal human myocardial-derived cells. The rationale for using neonatal cells for feasibility testing is based upon their more youthful biological phenotype of enhanced proliferation and survival, preservation of telomere length, and decreased level of senescence relative to cells derived from heart failure patients^14,15,19^.

hCCs created in the present study represent novel hybrid cells with phenotypic properties consistent with our previous observations of murine CCs. Having overcome the multiple aforementioned challenges of human myocardial-derived cell utilization, cell fusion is a feasible promising engineering approach to enhance functional properties of human cardiac cells and pave the way for ‘designer’ hCC exhibiting enhanced reparative properties.

## Methods

The authors declare that all supporting data are available within the article [and its online supplementary files]. The data that support the findings of this study are available from the corresponding author upon reasonable request.

### cCIC and MSC isolation

The study was designed in accordance with and approved by the institutional review committees at San Diego State University. Neonatal cells were derived from non-surgically obtained post-mortem cardiac tissue. Cells were isolated as previously published^12^. Briefly, heart samples were mechanically minced into 1mm^3^ pieces and digested in collagenase II (150 U mg/mL, Worthington, LS004174) followed by brief low speed (850 rpm, for 2 minutes) centrifugation to remove cardiomyocytes and tissue debris. The supernatant was subjected to magnetic activated cell sorting (MACS) using c-kit conjugated microbeads (Miltenyi Biotec, #130-091-332) and c-kit enriched cells were plated in human c-kit CIC media. The c-Kit negative population was further purified by MACS for MSC surface markers CD90 and CD105. c-kit CICs and MSCs were incubated at 37°C in 5% CO_2_ and used for cellular fusion between passages 5 and 10. Media used in the study are listed in Supplemental Table 1.

**Table 1.**
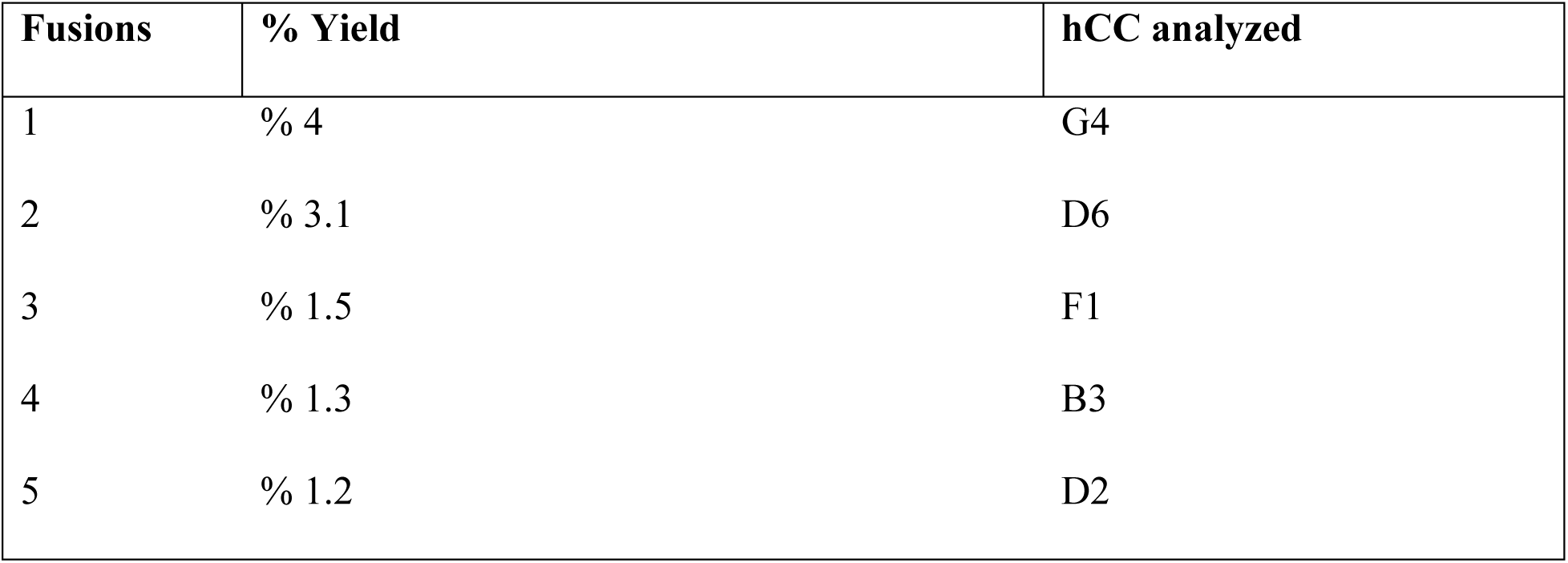
List of fusions.

| **Fusions** | **% Yield** | **hCC analyzed** |
| --- | --- | --- |
| 1 | % 4 | G4 |
| 2 | % 3.1 | D6 |
| 3 | % 1.5 | F1 |
| 4 | % 1.3 | B3 |
| 5 | % 1.2 | D2 |

### Lentiviral constructs and stem cell infection

Lentiviral plasmids and viral particles were created as previously described^8^. c-kit CICs and MSCs were stably transduced at passage 6 with PGK-EGFP-Puro and PGK-mCherry-Bleo respectively at a multiplicity of infection (MOI) of 40. Expression of EGFP and mCherry fluorescent proteins in c-kit CICs and MSCs was confirmed by fluorescent microscopy as well as flow cytometry. Lentivirally-modified cells were stabilized through multiple rounds of passaging and freeze/thaw cycles before fusion experiments.

### Cell fusion and creation of hCCs

Cell fusion was performed with GenomeONETM-CF EX Sendai Virus (Hemagglutinating Virus of Japan-Envelope or HVJ-E) Envelope Cell Fusion Kit (Cosmo Bio, USA) using the suspension method according to the manufacturer’s protocol. Briefly, cCIC-EGFP and MSC-mCherry populations were combined in 25µL of 1X cell fusion buffer at a ratio of 2:1 (total cell number of 100,000). The 2:1 ratio yielded the maximum number of double positive fused cells (hCCs) and therefore was chosen for the study. Inactivated Sendai virus was added to the cell mixture and incubated on ice for 5 minutes. Cell suspension was transferred to the 37°C incubator for 30 minutes with intermittent shaking every 5 minutes. Next, cells were plated in 2 ml fusion media on a well of a 6-well dish. Media was changed after 24 hours and cells were then cultured in fusion media for 4 days. Hybrid cells were identified and isolated by FACS sorting using parent cells in pure as well as mixed (but not fused) as controls. hCCs were plated on a 96-well microplate (1 cell per well) for clonal expansion. Clone nomenclature was based on well number in the plate. All hCCs were used between passages 4 to 6 for experiments.

### Light microscopy and measurement of cell morphology

Bright field images of hCCs and the parent cells were obtained using a LEICA DMIL microscope and cell boundaries were outlined by image analysis using ImageJ software. Cell surface area and length to width ratio were determined as previously described^20^.

### Proliferation and doubling time

c-kit CICs, MSCs and hCCs were plated on a 6-well dish at a density of 20,000 cells per well. At three time points (days 1,3, and 5) cells were trypsinized and counted manually using a hemocytometer. Cell proliferation rate was determined for each group as fold change over day 1. Cell doubling times were calculated using online population doubling time software (http://www.doubling-time.com/compute_more.php).

### Immunocytochemistry

c-kit CICs, MSCs and hCCs were plated on a two-well chamber slide (15,000 per well). Staining was performed following the protocol previously described^20^. Nuclei were stained with DAPI diluted in 1X PBS at room temperature for 15 minutes. Cells were imaged using a Leica TCS SP8 confocal microscope. Antibodies and dilutions are listed in Supplemental Table 2.

### Immunoblot analysis

c-kit CICs, MSCs and hCCs were plated on 100-mm plates. Protein cell lysates were collected using 200 µl of SDS–PAGE protein sample buffer. Proteins were separated on a 4-12% NuPage Novex Bis Tris gel (Invitrogen) and transferred onto a polyvinylidene fluoride (PVDF) membrane. Non-specific binding sites were blocked using Odyssey blocking buffer (LI-COR) and proteins were labeled with primary antibodies in 0.2% Tween in Odyssey blocking buffer overnight. After washing, blots were incubated with secondary antibodies in 0.2% Tween 20 in Odyssey blocking buffer for 1.5 hours at room temperature and scanned using the LICOR Odyssey CLx scanner. Quantification was performed using ImageJ software. Antibodies and their dilutions are listed in Supplemental Table 2.

### Real-Time PCR

c-kit CICs, MSCs and hCCs were plated on 100-mm plates. RNA lysate was collected from the culture and purified using Quick-RNA MiniPrep (Zymo Research). cDNA synthesis was conducted using iScript cDNA Synthesis Kit (Bio Rad). qPCR was performed on a Bio-Rad CFX real time cycler using iQ SYBR Green (Bio Rad) and gene specific primers. Signals were normalized to 18S for analysis. Data were calculated using the ΔΔC(t) method. Primers are listed in Supplemental Table 3.

### Ploidy and cytogenic analysis

c-kit CICs, MSCs and hCCs were plated on a 6-well plate at a density of 50,000 cells per well. Cells were collected the following day and centrifuged at 1200 rpm for 5 minutes, the pellet was re-suspended in 70% ethanol, and stored at −20°C for at least 24 hours prior to use. After centrifugation at 1300 rpm for 5 minutes, cell pellet was re-suspended in 300 ul of Propidium Iodide incubated at 37°C for 15 minutes prior to flow cytometry analysis. Cytogenetic analysis of c-kit CICs, MSCs, and hCCs (G4) plated at a density of 300,000 cells on 2500 mm^2^ flasks was performed by KaryoLogic, Inc (North Carolina; www.karyologic.com).

### Cell death assay

For reactive oxygen injury, c-kit CICs, MSCs and hCCs were plated on a 6-well plate at a density of 60,000 cells per well. Cells were subjected to low serum media for 24 hours (depleted to 25% of growth media serum level) followed by 4 hours of hydrogen peroxide (350 μmol/L) treatment. Annexin V and Sytox Blue staining was performed to label apoptotic and necrotic cells and cell death was measured using FACS Aria (BD Biosciences).

For ischemia-reperfusion injury, cCICs, MSCs and hCCs were seeded on 6-well plates at a density of 60,000 cells per well. The following day, media was replaced with Kreb’s Heinsleit buffer (KH buffer) to induce glucose starvation, and cells were transferred to a hypoxic incubator with 1% oxygen tension for 3 hours to simulate ischemia. Cells were re-exposed to regular growth media and incubated in a standard cell culture incubator with ambient (21%) oxygen for 24 hours to simulate reperfusion. Annexin V and Sytox Blue staining was performed to label apoptotic and necrotic cells and cell death was measured using FACS Aria (BD Biosciences). Cells cultured in growth media in normoxic conditions and cells subjected to Kreb’s Heinsleit buffer (KH buffer) in hypoxic condition were used as the controls of the experiment to measure basal and hypoxia-induced cell death, respectively. KH buffer and the respective media used inside the hypoxic glove box were equilibrated in hypoxia overnight before starting the experiment.

### NRCM co-culture with stem cells

NRCMs were isolated as previously described^21,22^ and seeded in a 6-well dish at a density of 200,000 per well in M199 media with 15% fetal bovine serum (FBS). The following day, cells were incubated in media with 10% FBS for 8 hours followed by 24-hour serum depletion in serum free media. Stem cells (cCICs, MSCs, combination of cCICs and MSCs, hCCs) were added to the culture at a 1:5 ratio. The slow growing clone B3 was excluded from this experiment due to a low expansion rate. After 24 hours in co-culture, cells were stained with Annexin V and Sytox Blue. Unlike CardioChimeras or their parent cells, the NRCMs were non-transduced allowing for separation by fluorescent activated cell sorting (FACS) of negative cells (NRCMs) versus GFP+, mCherry+ or GFP+/mcherry+ cells. Thus, parental and CardioChimera cells were removed from the population for survival analysis of NRCMs, which was completed by flow cytometry using the FACS Aria. Controls for the NRCMs included 1) culture in serum free media alone (SF), 2) “rescue” by replenishment with M199 media + 10% serum, or 3) constant culture in M199 media + 10% serum for the duration of the experiment.

### Statistical analysis

All data are expressed as mean ±SEM. Statistical significance between two groups was assessed using two-tailed t-test, or one-way ANOVA or two-way ANOVA for multiple comparisons, with the Dunnett and Tukey tests as post hoc tests to compare groups to a control group in Graph Pad Prism v5.0. or Microsoft excel. P<0.05 was considered statistically significant.

## Results

### hCCs creation from c-kit CIC and MSC cell fusion

Human c-kit CICs expressing GFP and human MSCs expressing mCherry, incubated in the presence of inactivated RNA Sendai virus, underwent cellular fusion to form mononuclear hybrid cells^23,24^ (Figure 1A). A group of mixed c-kit CICs and MSCs without the viral treatment was used as the negative control for the fusion experiment (Supplemental Figure 1E). Readily detectable uniform fluorescence labeling of parental c-kit CICs and MSCs is required for optimal yield of double fluorescent hybrids. Flow cytometry analysis shows 96.6% of c-kit CICs and 92% of MSCs expressed EGFP and mCherry, respectively (Supplemental Figure 1A-D). Double fluorescent hybrids were sorted into 96-well microplates for clonal expansion to derive hybrid clones called hCCs. Five unique hCCs named D6, F1, G4, D2 and B3 were derived from five independent fusion experiments with 1-4% efficiency (Table 1). Dual fluorescent positivity of hCCs was validated by fluorescence microscopy (Supplemental Figure 2). Immunocytochemistry with respective EGFP and mCherry antibodies confirmed integration of genomic content from both parent cells into hCC that was maintained after clonal expansion and passaging in culture (Figure 1B). Parent cells immunolabeled for EGFP and mCherry exhibited appropriate single wavelength fluorescence signal, validating antibody specificity for their respective fluorophores (Supplemental Figure 3). Immunoblotting results corroborate flow cytometry and microscopy observations for expression of both fluorescent marker proteins (Figure 1C). Collectively, these findings indicate successful cellular fusion and stability of the chimeric state after clonal expansion.

**Figure 1.**
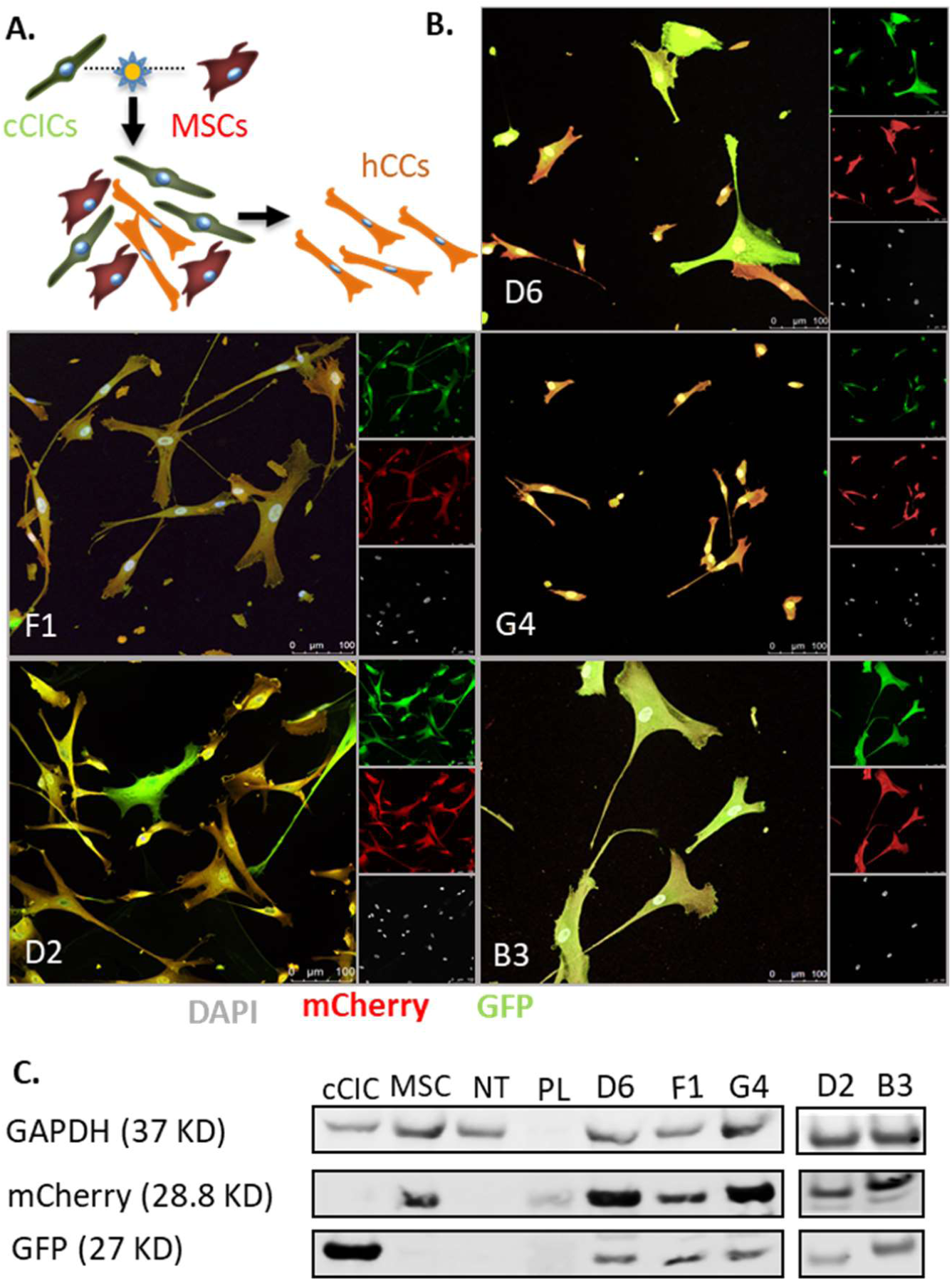
hCCs are created from cCIC and MSC cell fusion. A. Schematic of the cell fusion. Phase A: fusion-suspension of cCICs and MSCs. Phase B: initial fused cells undergo mitotic event to combine chromatin content. Phase C: Sorting of successfully fused cell. **B**. ICC of the hCCs for GFP and mCherry. **C**. Immunoblot analysis of cCIC-GFP, MSC-mCherry, Non-Transduced (NT) MSC, and hCCs. PL: protein ladder. GAPDH is used as the loading control.

**Figure 2.**
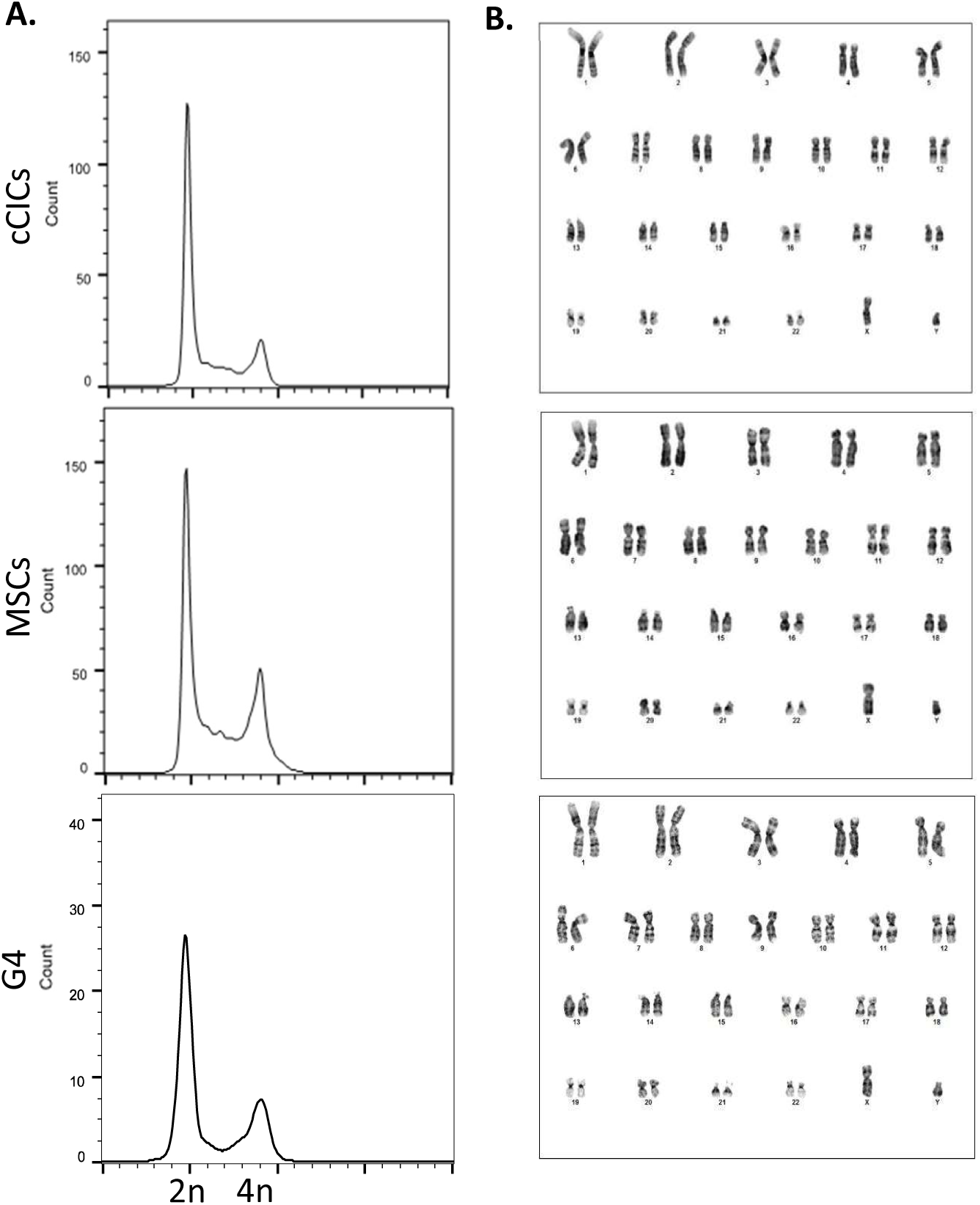
hCCs reveal ploidy status and DNA content corresponding to the parent cells. **A**. Flow Cytometry plots for PI/RNAse staining of parent cells and hCCs, G4. N=3 independent experiments. **B**. Cytogenetic analysis of parent cells and hCC, G4.

**Figure 3.**
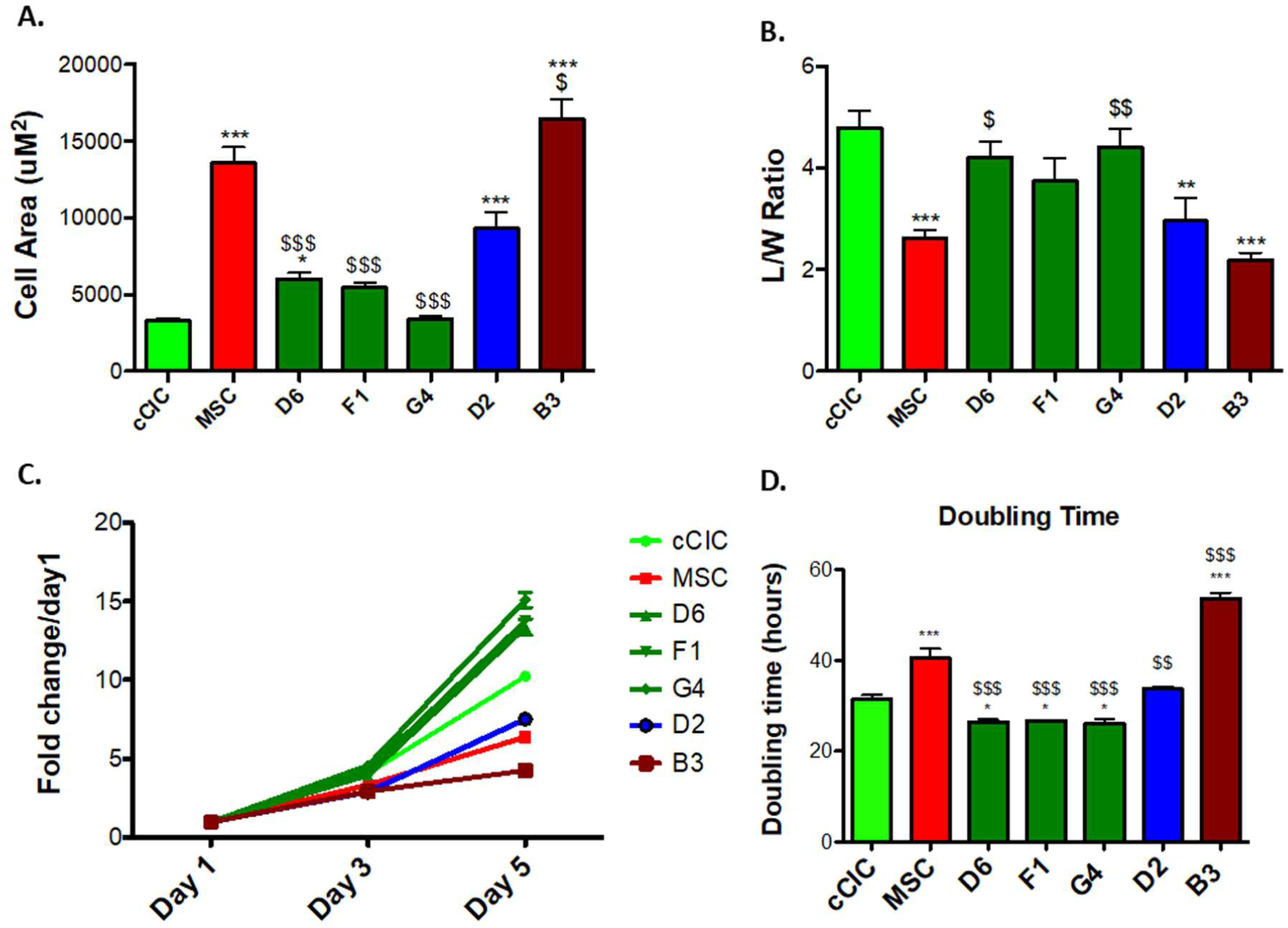
Morphometric and proliferative characteristics of hCCs. **A**. Cell surface area and **B**. length to width ratio of the parent cells and hCCs. **C**. Proliferation rate of parent cells and hCCs represented as fold change over day 1, and **D**. cell doubling time represented in hours. N=3 independent experiments. Statistical analysis was performed by One-Way ANOVA multiple comparison with Dunnett. *P<0.05 Vs cCIC, **P<0.001 Vs cCIC, ***P< 0.0001 Vs cCIC, $P<0.05 Vs MSC, $$P<0.001 Vs MSC, $$$P<0.0001 Vs MSC. Error bars are ± SEM.

### hCCs possess diploid DNA content

Fusion of two parental lines resulted in elevated ploidy in murine CCs DNA content^8^. In comparison, hCCs possess diploid (2n) DNA content comparable to the parent cells as assessed using flow cytometry analysis (Figure 2A and Supplemental Figure 4). Since all hCCs exhibited 2n ploidy status, a representative line (hCC G4) was chosen for karyotypic analysis that confirmed normal male karyotype with 46 chromosomes including XY as observed in G-banded spreads (Figure 2B). Therefore, hCCs maintain normal 2n ploidy content relative to the parent cells, in contrast to increased ploidy of murine CCs^8^.

**Figure 4.**
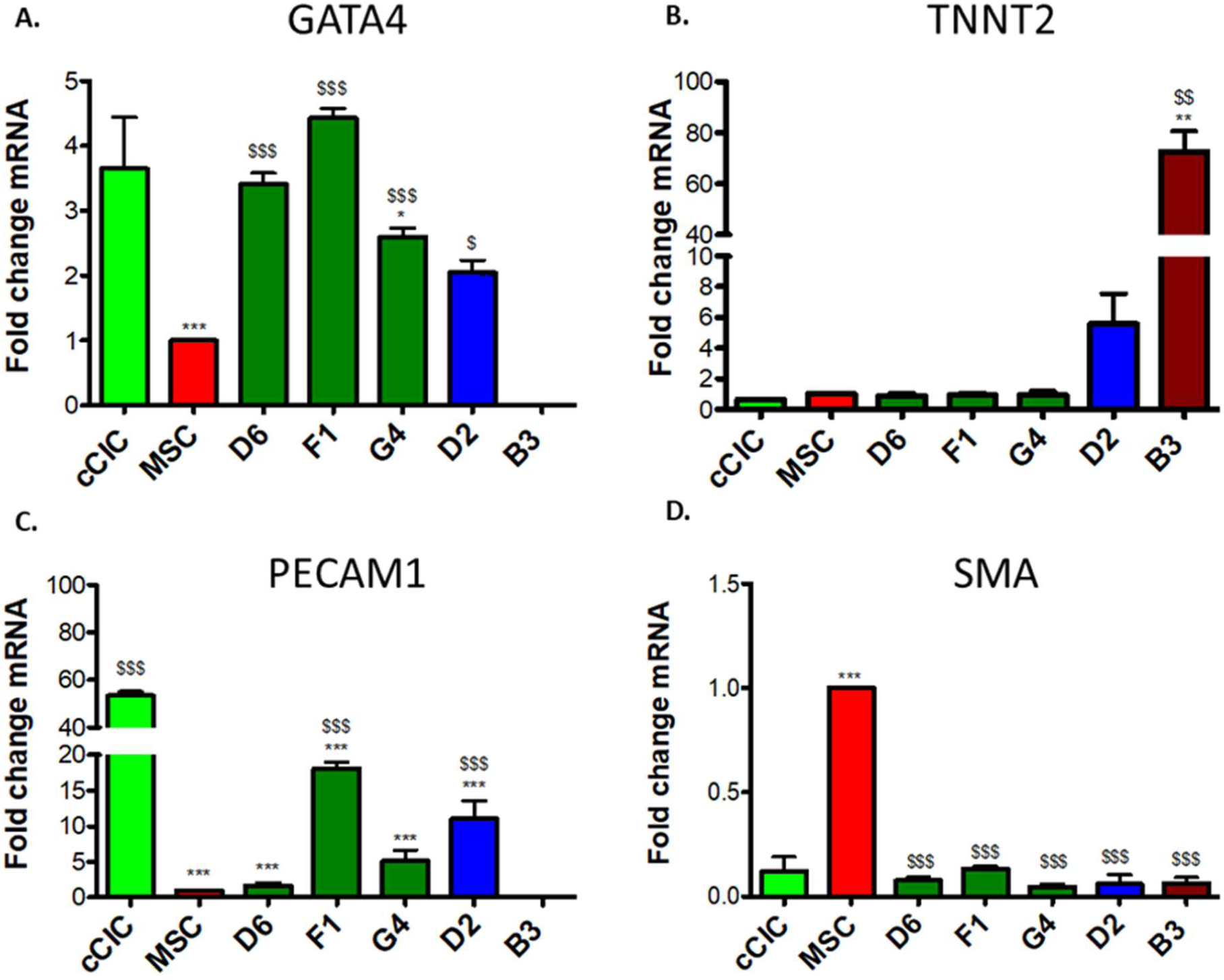
hCCs exhibit variable expression level of cardiomyogenic commitment markers. mRNA expression level of **A**. GATA4, **B**. TNNT2, **C**. PECAM1, and **D**. SMA for parent cells and hCCs. N=3 independent experiments, 3 replicates per group per experiment. All genes expression is normalized to ribosomal 18s and represented as a fold change relative to MSCs. Statistical analysis was performed by One-Way ANOVA multiple comparison with Dunnett. *P<0.05 Vs cCIC, **P<0.001 Vs cCIC, ***P< 0.0001 Vs cCIC, $P<0.05 Vs MSC, $ $P<0.001 Vs MSC, $ $ $P<0.0001 Vs MSC. Error bars are ± SEM.

### hCCs phenotypic and proliferative characteristics resemble parent cells

Murine CCs characteristics of survival and proliferation were comparable to parental lines albeit with increased DNA content^8^. Cellular properties of select hCCs were assessed for phenotypic traits of morphology and growth rate relative to parental lines, chosen to represent typical examples for variability of hCC characteristics. Cellular morphology measured using bright field images of the hCCs and parent cells reveals a range of cell surface area, length to width ratio (Figure 3). hCCs D6, F1, and G4 exhibited cell surface area similar to cCICs that was significantly lower than MSCs (Figure 3A). Length to width ratio of these clones was significantly greater compared to MSCs and was comparable to that of cCICs (Figure 3B). Proliferation rate of these hCCs was increased relative to parental cells with a doubling time of approximately 24 hours and were therefore characterized as fast growth (Figure 3C,D). Cell surface area of hCC D2 was significantly greater than the cCICs and similar to the MSCs (Figure 3A). Length to width ratio for hCC D2 was significantly decreased relative to cCICs, instead resembling that of MSCs (Figure 3B). Proliferation rate of hCC D2 was intermediate compared to parent cells with a doubling time of 32 hours and was identified as a medium growth (Figure 3C,D). Lastly, hCC B3 cell surface area was increased, length to width ratio was reduced, and proliferation rate was slow compared to either of the parent cells, resulting in designation of slow growth (Figure 3A-D). Collectively, these data profile phenotypic properties of hCCs with cell surface areas and length to width ratios that correspond to their proliferative rates, consistent with previous findings for murine CCs^8^.

### Profiling for commitment and secretory gene expression reveals hCC variability

Gene expression profiling revealed significant heterogeneity between various murine CCs^8^, so commitment and secretory gene profiles for hCCs were assessed to find if similar diversity was present. Cardiac lineage markers including cardiac type troponin T2 (TNNT2), GATA Binding Protein 4 (GATA4), Platelet and Endothelial Cell Adhesion Molecule 1 (PECAM1), and smooth muscle actin (SMA) were investigated by measuring mRNA expression level by qPCR analysis. Early cardiac transcription factor GATA4 was significantly upregulated in all hCCs except B3 in which GATA4 expression was not detected when compared to MSCs. When measured relative to cCICs, GATA4 was approximately one-fold higher in F1, unchanged in D6 and G4, and downregulated in D2 (Figure 4A). Cardiomyocyte marker TNNT2 level was reduced in fast growing hCCs D6, F1 and G4 as opposed to medium and slow growing clones D2 and B3 (Figure 4B). PECAM1 level was higher in F1, G4 and D2 clones than in MSCs, but lower than cCICs. PECAM1 was not detected in B3 (Figure 4C). All hCCs significantly downregulated SMA relative to MSCs resembling closely to cCICs (Figure 4D). Collectively, these data demonstrate that fast-growing hCCs upregulate expression of early cardiac transcription factor GATA4 and downregulate expression of maturational cardiac marker TNNT2, reflective of their cardiac progenitor-like phenotype. In contrast, medium and slow growing hCCs exhibit a more committed mRNA profile from this cursory assessment with select transcripts.

hCCs were also analyzed for expression of pro-survival paracrine factors that mediate protective effects. HB-EGF, a secreted glycoprotein involved in wound healing and cardiac development^25,26^, was significantly upregulated in hCC F1. In comparison, hCCs D6, G4 and D2 expressed HB-EGF at levels intermediate between the two parent cell ranges. HB-EGF expression was undetectable in hCC B3 clone (Figure 5A). HGF, a paracrine growth, motility and morphogenic factor^27,28^ was expressed at low levels in hCC D6, F1, G4 and D2, but at a comparable level to MSCs in B3 (Figure 5B). hCCs F1 and D2 expressed high levels of chemotactic factor SDF relative to cCICs but lower than in MSCs. SDF expression was downregulated in D6 and G4 and undetectable in B3 (Figure 5C). FGF2, an important growth factor for wound healing, angiogenesis and cellular proliferation^29^ was highly upregulated in B3. hCC F1 and D2 increased FGF2 expression relative to cCICs and were more reminiscent of MSCs, while hCC D6 and G4 expressed lower levels of FGF2 compared to both parent cells (Figure 5D). In summary, similar to what was previously observed with murine CC^8^, profiling of hCC reveals distinct transcriptome profiling signatures between the clones.

**Figure 5.**
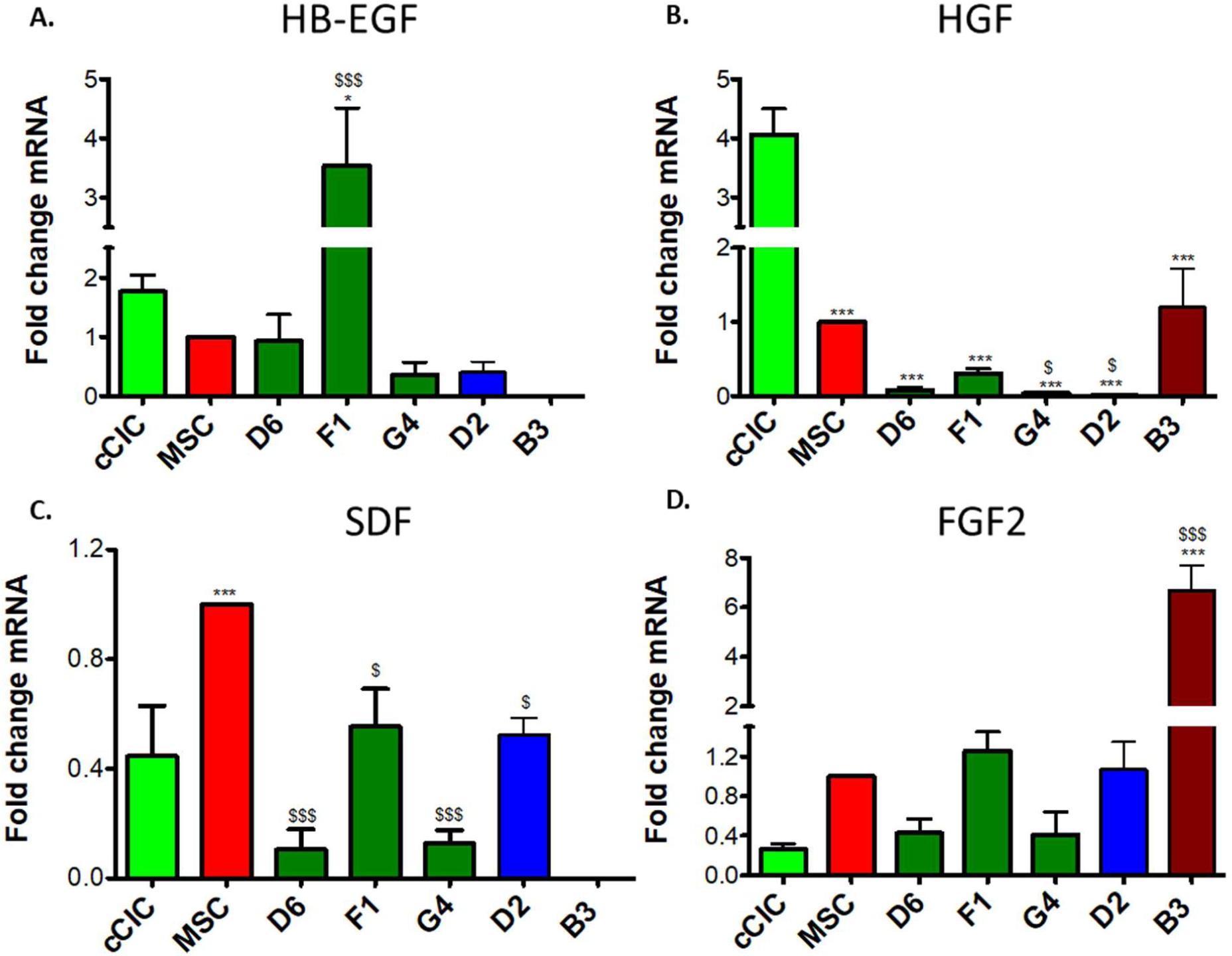
hCCs exhibit variable secretory gene profiles. mRNA expression level of **A**. HB-EGF, **B**. HGF, **C**. SDF, and **D**. FGF2. N=3 independent experiments, 3 replicates per group per experiment. All genes expression is normalized to ribosomal 18s and represented as a fold change relative to MSCs. Statistical analysis was performed by One-Way ANOVA multiple comparison with Dunnett. *P<0.05 Vs cCIC, **P<0.001 Vs cCIC, ***P< 0.0001 Vs cCIC, $P<0.05 Vs MSC, $ $P<0.001 Vs MSC, $ $ $P<0.0001 Vs MSC. Error bars are ± SEM.

### Survival in response to environmental stress is enhanced in hCC

hCCs response to stress was assessed by serum deprivation with cell death quantitation by flow cytometric measurement of necrosis and apoptosis. Cell death rates were similar between hCCs and MSCs but lower than in cCICs (Figure 6A). Response to oxidative stress induced by hydrogen peroxide treatment (350 µmol/L) in serum free conditions reveals significantly lower rates of apoptosis and necrosis compared to either parental line with correspondingly higher rate of survival (Figure 6B), consistently present even when media formulations for parental lines (cCIC or MSC) are used for hCC culture (Supplemental Figure 5; D2 line). Simulated ischemia-reperfusion injury was also used as an environmental stress with cells subjected to KH buffer in hypoxia, and growth media in normoxia as experimental controls. Lower necrotic cell death was found for all hCCs compared to parent cells. Similar apoptosis rates were observed comparing hCCs to the parental lines with the exception of D2, which exhibited higher apoptotic cell death (Figure 6C). Correspondingly, survival of all the hCCs except D2 was not significantly different from either parental cell. Similar percentages of apoptotic, necrotic and live cells were observed in experimental controls (Supplemental Figure 6). In conclusion, unlike susceptibility to environmental stress in murine CCs similar to parent lines^8^, hCCs exhibit enhanced resistance to oxidative stress.

**Figure 6.**
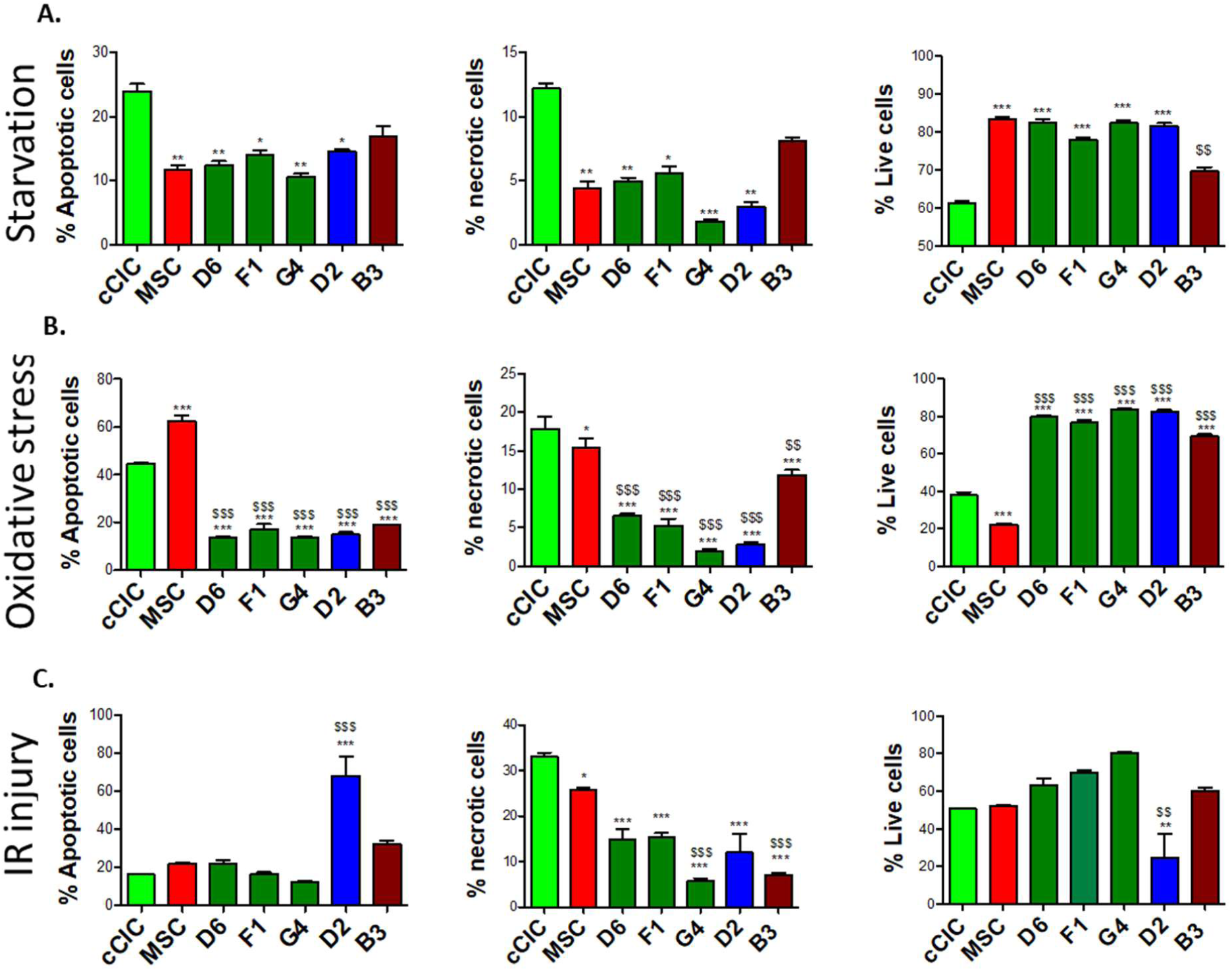
hCCs demonstrate enhanced survival in response to stress. Percentage of apoptotic, necrotic and live cells following **A**. serum starvation, **B**. oxidative stress, and **C**. IR injury. N=3 independent experiments. Statistical analysis was performed by Two-Way ANOVA multiple comparison with Tukey. *P<0.05 Vs cCIC, **P<0.001 Vs cCIC, ***P< 0.0001 Vs cCIC, $P<0.05 Vs MSC, $$P<0.001 Vs MSC, $$$P<0.0001 Vs MSC. Error bars are ± SEM.

### Cardiomyocyte survival is enhanced by hCC co-culture

Protective effects in response to serum deprivation challenge of NRCMs in vitro is similar between murine CCs or their corresponding parental cells^8^. The protective effect of hCCs was similarly assessed, with serum deprivation challenge of NRCMs that prompted significantly increased cell death which was partially mitigated by addition of serum for 24 hours (Figure 7). Clone B3 was excluded from further experiments due to slow expansion rate, undergoing replicative senescence after passage 10 in culture. In comparison, lines D6, F1and G4 expansion rate slowed after passage 10, yet had not reached replicative senescence. Line D2 maintained consistent proliferation rate for more than 10 passages, currently expanded more than 27 passages without arrest (Supplemental Figure 7). NRCMs showed improved survival mediated by each hCC in response to serum deprivation, co-culture with hCC D2 yielding the highest rate of survival (Figure 7). NRCMs co-cultured with cCICs exhibited high level of survival in response to serum deprivation. Co-culture of MSCs and hCCs G4 partially rescued NRCMs resembling the effect of serum addition for 24 hours. hCCs D6 and F1 exerted intermediate protective effect on NRCMs compared to parental cells while hCCs D2 provided the highest protective effect on NRCMs, exceeding even that of NRCMs in high serum. Except for hCC G4 which exhibited protective effect similar to MSCs, most hCCs resembled cCICs in co-culture effect upon NRCMs. Therefore, in contrast to murine CCs with protective effect similar to their corresponding parent cells, hCCs exhibit individually distinct variability in protective action.

**Figure 7.**
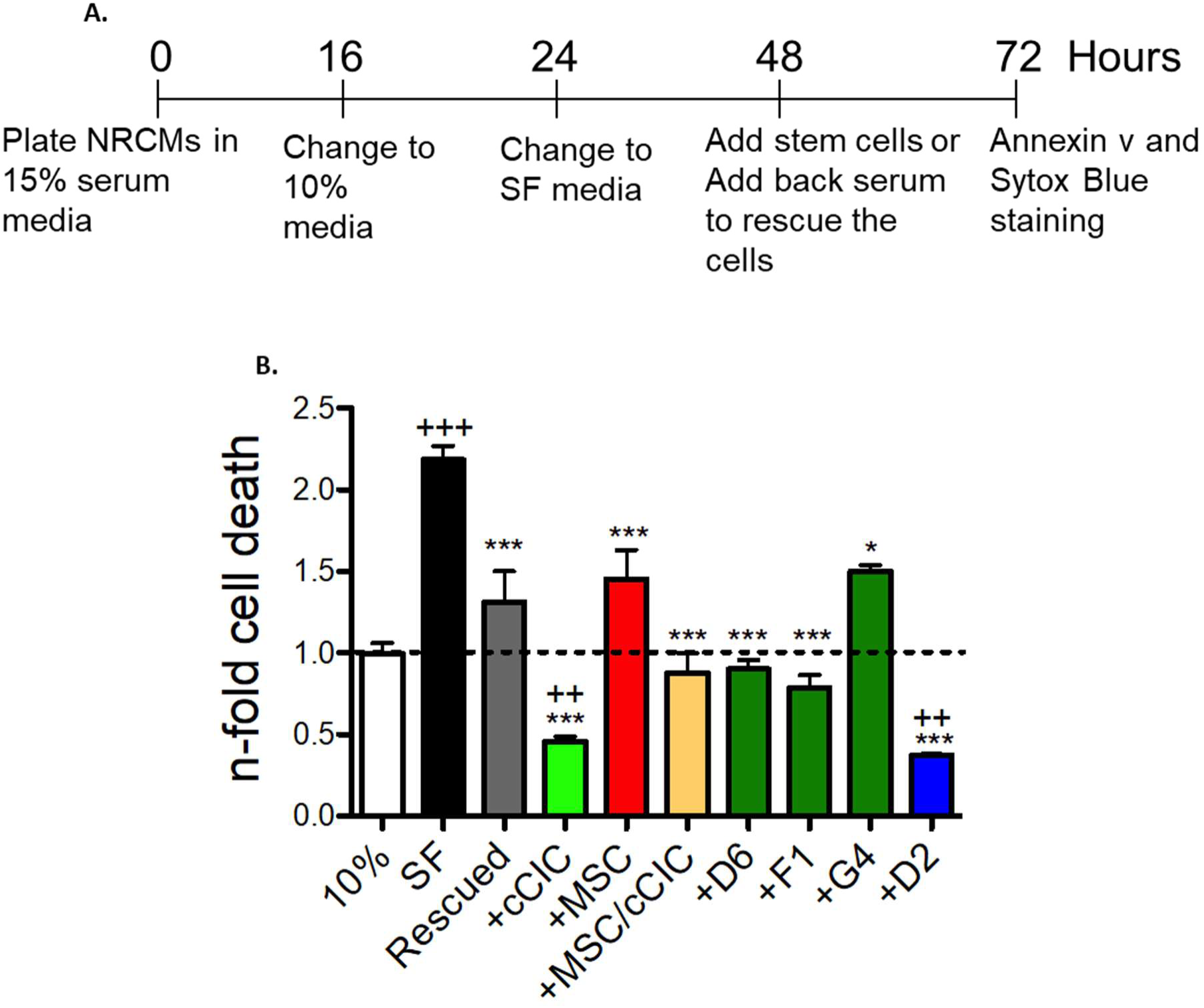
hCCs promote cardiomyocytes survival in response to serum deprivation. **A**. NRCM co-culture protocol. **B**. Cardiomyocytes cell death in 10% serum media (10%), serum free media (SF), rescued and in presence of cCICs, MSCs, combinatorial group of MSCs and cCICs (+MSCs/cCICs), D6, F1, G4 and D2. Values presented are fold change of Annexin v^+^ Sytox Blue^+^ cells relative to 10% serum media (also represented as dashed line, 1.0). N=3 independent experiments. Statistical analysis was performed by One-Way ANOVA multiple comparison with Dunnett. *P<0.05 Vs SF, ***P< 0.001 Vs SF, ++ P<0.01 Vs 10%, +++ P<0.001 Vs 10%.

## Discussion

Myocardial repair and regeneration research continues to progress on multiple fronts supported by over a decade of cumulative studies despite a recent “whipsaw” movement of controversies related to the terminology and biology of “cardiac stem cells”^30,31^. Although cardiogenic potential of “cardiac stem cells” remains debatable^32,33^, cardioprotective effects have been repeatedly demonstrated by our group and others^13,34,35,36^ with “cardiac stem cells”, now referred to as cCIC in this report to avoid being misconstrued as cardiogenic cells. Furthermore, cardioprotective effects of cCICs are enhanced by combinatorial delivery with MSCs^6^. These findings are encouraging, but efficacy of such adoptive transfer cell therapies is inherently limited by multiple factors including poor cell survival, engraftment, and persistence that can be overcome to some extent by ex vivo cell engineering. Our group has focused upon potentiation of myocardial repair through use of modified cells created by genetic engineering^9,10,14,37^, fusion^8^, and combinatorial culture / delivery^38^ (Megan Monsanto, unpublished data, [2019]). Implementation of these ‘unnatural’ approaches to create engineered cell types and combinations is deliberately intended and rationally designed to overcome inherent limitations of normal mammalian myocardial that lacks endogenous cellular reservoirs to repair and restore myocardial structure and function following pathological damage. The rationale for CC in our initial study was to avoid issues related to variable survival, engraftment, persistence, and proliferation rates of cCIC versus MSC when mixing these two distinct populations together for cell therapy administration. We posited that creating a novel fused hybrid CC would deliver significantly enhanced reparative activity compared to each parental cell type used either alone or in mixed single cell suspensions; a postulate that was validated in a syngeneic murine infarction model^8^. The next logical advance of this patented technology (US Patent #20160346330A1) was to apply a comparable approach in the context of human cCIC and MSC. Species-specific characteristics between murine versus human biology result in differences in the creation and phenotypic properties of CC, as reinforced by results in this report.

Comparison between human versus murine CC reveal both similarities as well as important distinctions. Specifically with respect to similarities between mouse CC and hCC, there were a few notable parallels including 1) variable proliferative characteristics (slow, medium, fast), 2) variable CC size, 3) comparable PECAM 1 expression (intermediate level in both mouse and human CCs compared to the parent cells), and 4) cardiac troponin T expression at high level relative to parental lines in hCC D2 and B3 (medium and slow grower lines, respectively) similar to mouse CC versus parental lines. In comparison, differences between mouse CC and hCC were numerous and significant as demonstrated by 1) higher survival level compared to parental cells in response to oxidative stress by hCCs, whereas murine CC survival was either similar or lower than parental cells, 2) normal diploid DNA content of hCC compared to tetraploidy in murine CC, 3) SMA expression was lower in hCC than either parental cell, but higher in mouse CCs than the cCIC parental cell, 4) cardiac troponin T expression level in hCC similar to fast growing parental lines versus higher level expression in mouse CC, and 5) variable protective effects in NRCM co-culture studies of hCC relative to parental cells versus similar protective effects between murine CC and parental cells. Even with an admittedly small sampling size of two murine CCs and 5 hCCs there are demonstrable biological and phenotypic differences between species-specific CC warranting further characterization.

Polyploidy observed in mouse CC was notably absent from human CC lines. The diploid (2n) DNA content of hCC (Figure 2) contrasts with that of murine CCs which had increased DNA content^8^. Cytogenic analysis of hCC G4 further confirmed a normal karyotype with 46 chromosomes. The most straightforward interpretation of this finding is that human cell fusions do not tolerate polyploidy as a permissive state for mitotic growth, unlike their murine counterparts. hCCs may acquire one copy of each chromosome complement from either of the parent cells leading to no change in ploidy content, despite being successfully fused. This cytogenetic state eliminates concerns regarding fusion-induced increased DNA content and aneuploidy which can lead to genomic instability and cellular senescence^39^. All hCCs retained cellular properties corresponding to cCIC and MSCs consistent with chromosomal mosaicism in somatic cells without significantly altering stem cell behavior^40^.

MSCs mediate cell survival through secretion of pro-survival chemokines and cytokines including SDF-1 and IGF-1^41,42^. hCCs expressed growth factors and cardioprotective cytokines including EGF and SDF and were resistant against starvation-induced cell death (Figure 5A,C and Figure 6A). Similarly, hCCs demonstrated enhanced survival in response to oxidative stress (Figure 6B) and attenuated necrotic cell death in response to I/R injury (Figure 6B,C). When co-cultured with NRCMs, hCC G4 exhibited cardioprotective effect similar to MSCs whereas hCCs F1, D6 and D2 exhibited cardioprotective effect similar to cCIC with hCC D2 exhibiting the highest cardioprotective activity (Figure 7). Cellular cross talk survival signals might be facilitated in CC, since exosome contents such as MSC-secreted micro-RNA 21 protects c-Kit^+^ stem cells from oxidative injury through the PTEN/PI3K/Akt signaling pathway^43^. Paracrine signaling activity has been suggested as the mechanistic basis for cardioprotection using cell therapy^44,45,46^, so the relevance of the secretome from hCCs deserves further detailed analyses to establish a putative role in enhancing survival signaling. The potential contribution of cell fusion to heart regeneration is essentially unexplored despite the established role of cell fusion as a reprogramming mechanism leading to enhanced proliferation and growth rate in differentiated cells^47^. The findings presented here suggest cell fusion as a reliable and stable technique to generate human hybrid cells. cCICs and MSCs are ideal cellular candidates for cell fusion. They can be consistently isolated and expanded in laboratory settings and they are well established cell candidates for cellular therapy based on the results of basic biological studies and clinical trials^4,48^. In the present study, cCICs and MSCs were used at a ratio of 2:1 but cell ratios used for fusion can influence the fate and functional potential of resultant hybrids. Therefore, modifying the cell number ratio prior to fusion could be a promising future direction leading to the design of hybrids with individualized phenotypic characteristics. Moreover, adapting cell fusion ratios for future studies may promote creation of hCCs with more consistent biological profiles.

Cell fusion is a tractable engineering approach to generate a novel cell population incorporating characteristics of two different human cardiac derived interstitial cell populations into a single cell. Hybrid CCs possess molecular and phenotypic characteristics similar to previously established murine CCs with enhanced myocardial repair potential^8^. However, cell engraftment, cardiomyogenic and reparative potential of hCCs need to be determined and are the subject of the future investigation that could support next steps toward translational application of hCCs.

## Acknowledgments

F. Firouzi, S.S. Choudhury and M.A. Sussman designed experiments. F. Firouzi, S.S. Choudhury, K. Broughton and A. Salazar performed experiments and analyzed data. F. Firouzi and M.A. Sussman wrote the manuscript. All the authors read and approved the article.

We gratefully acknowledge Dr. Christopher Glembotski and Erik Blackwood for providing the Sussman lab with NRCMs used for co-culture experiment as well as Dr. Roland Wolkowicz, Cameron Smurthwaite and Ruby Cotton for their assistance with cell sorting and flow cytometry analysis. We thank Dr. Barbara Bailey from SDSU Mathematics department for her help with the statistical analysis of the results. All persons named in the Acknowledgments section have provided the corresponding author with permission to be named in the manuscript.

## Non-standard Abbreviations and Acronyms

BDM: 2,3-Butanedione 2-monoxime
CC: CardioChimera
cDNA: Complementary DNA
DAPI: 4’,6-diamidino-2-phenylindole
DMEM: Dulbecco’s modified eagle medium
ES FBS: Embryonic stem cell FBS qualified
FACS: Fluorescence-Activated Cell Sorting
FBS: Fetal bovine serum
FGF2: Basic fibroblast growth factor
GAPDH: Glyceraldehyde 3-phosphate dehydrogenase
GATA4: GATA binding protein 4
GFP: Green fluorescent protein
HB-EGF: Heparin binding EGF like growth factor
HBSS: Hank’s balanced salt solution
hCC: Human CardioChimera
hcCICs: Human **c-kit+** cardiac interstitial cells (previous nomenclature of cardiac progenitor cells has been updated due to controversies in the field)
HEPES: 4-(2-hydroxyethyl)-1-piperazineethanesulfonic acid
HGF: Hepatocyte growth factor
hMSC: Human mesenchymal stem cell
IL-6: Interlukin-6
I/R: Ischemia-reperfusion
LIF: Leukemia inhibitory factor
MI: Myocardial infarction
NRCM: Neonatal rat cardiomyocyte
PBS: Phosphate buffered saline
PECAM1: Platelet endothelial cell adhesion molecule or CD31
PI: Propidium iodide
SDF: Stromal derived factor
SMA: Smooth muscle actin
TNNT2: troponin T2, cardiac type

## Source of Finding

M.A. Sussman is supported by NIH grants: R01HL067245, R37HL091102, R01HL105759, R01HL113647, R01HL117163, P01HL085577, and R01HL122525, as well as an award from the Foundation Leducq. F.Firouzi is supported by Grace Hoover Sciences Scholarship. K.M. Broughton is supported by NIH grant F32HL136196.

