## Supplemental data for "Human CardioChimeras: Creation of a Novel ‘Next Generation’ Cardiac Cell"

Short title: **Human CardioChimera: Next Generation Cardiac Cell**

**Corresponding Author:

Mark A. Sussman, PhD

SDSU Heart Institute and Department of Biology

San Diego State University, 5500 Campanile Drive

San Diego, CA 92182

**Subject codes:** Basic Science Research; Cell Therapy; Myocardial Biology; Stem Cells; Cell Biology/Structural Biology

**SUPPLEMENTAL MATERIAL**

**Supplemental Tables**

**Supplemental Table 1. List of media**

| **Media** | **Components** |
|---|---|
| Human cCIC Media | 10% ES FBS, 1% Penicillin-Streptomycin-Glutamine (100X), 5 mU/mL human erythropoietin, 10 ng/mL human recombinant basic FGF, 0.2 mmol/L L-Glutathione in F12 HAM's (1x) |
| Human MSC Media | 20% FBS, 1% Penicillin-Streptomycin-Glutamine (100X) in 10.1 g/L Minimum Essential Medium Eagle, Alpha Modification |
| Human Fusin Media | 15% ES-FBS, 1% Penicillin-Streptomycin-Glutamine (100X), 5 mU/mL human erythropoietin, 10 ng/mL human recombinant basic FGF, 0.2 mmol/L L-Glutathione, 0.2 mg/ml human IL-6, 0.25 mg/ml human LIF in F12 HAM's (1x) |
| KH Buffer | 125 mmol/L NACL, 8 mmol/L KCL, 1.2 n mol/L $KH_2PO_4$, 1.25 m mol/L $mgSO_4$, 1.2 m mol/L $CaCl_2$, 6.25 m mol/L $NaHCO_3$, 20 m mol/L deoxy-glucose, 5 m mol/L Na-lactate, 20 m mol/L HEPS |
| NRCM Plating Media | 15% FBS, 1% Penicillin-Streptomycin-Glutamine (100X) in medium 199 |
| NRCM Maintenance Media | 10% FBS, 1% Penicillin-Streptomycin-Glutamine (100X) in medium 199 |

**Supplemental Table 2. List of Antibodies**

| Antibody | Manufacturer | # Catalog | Dilution | Application |
|---|---|---|---|---|
| GFP anti rabbit | Thermofisher | A11122 | 1:80 | Immunocytochemistry |
| mCherry anti Rat | Thermofisher | M11217 | 1:80 | Immunocytochemistry |
| DAPI | Sigma | D9542 | 1:5000 | Immunocytochemistry |
| GFP anti Goat | Rockland | 35059 | 1:500 | Immunoblotting |
| mCherry anti mouse | Abcam | Ab125096 | 1:500 | Immunoblotting |
| GAPDH anti Goat | Sicgen | AB0067 | 1:3000 | Immunoblotting |
| Annexin V-APC | Biosciences | 550475 | 1:175 | Cell death assay |
| Sytox Blue | Life Technologies | S11348 | 1:2000 | Cell death assay |
| Propidium Iodide | Invitrogen | P3566 | 1:75 | Ploidy analysis |

**Supplemental Table 3. qRT PCR primer list**

| mRNA Primers | Forward | Reverse |
|---|---|---|
| GATA 4 | CTCAGAAGGCAGAGAGTGTGTCAA | CACAGATAGTGACCCGTCCCAT |
| TNNT2 | GGAGAGAGAGTGGACTTTATG | CCTCCTCTTTCTTCCTGTTTC |
| PECAM1 | CCAAGCCCGAACTGGAATCT | CACTGTCCGACTTTGAGGCT |
| SMA | CCCAGCCAAGCACTGTCAGGAATCCT | TCACACACCAAGGCAGTGCTGTCC |
| HB-EGF | ACAAGGAGGAGCACGGGAAAAG | CGATGACCAGCAGACAGACAGATG |

| | | |
|---|---|---|
| HGF | GGCTGGGGCTACACTGGATTG | CCACCATAATCCCCCTCACAT |
| SDF | CAGTCAACCTGGGCAAAGCC | AGCTTTGGTCCTGAGAGTCC |
| FGF2 | CTGGCTATGAAGGAAGATGGA | TGCCCAGTTCGTTTCAGTG |
| 18s | CGAGCCGCCTGGATACC | CATGGCCTCAGTTCCGAAAA |

### Supplemental Figures and Figure legends

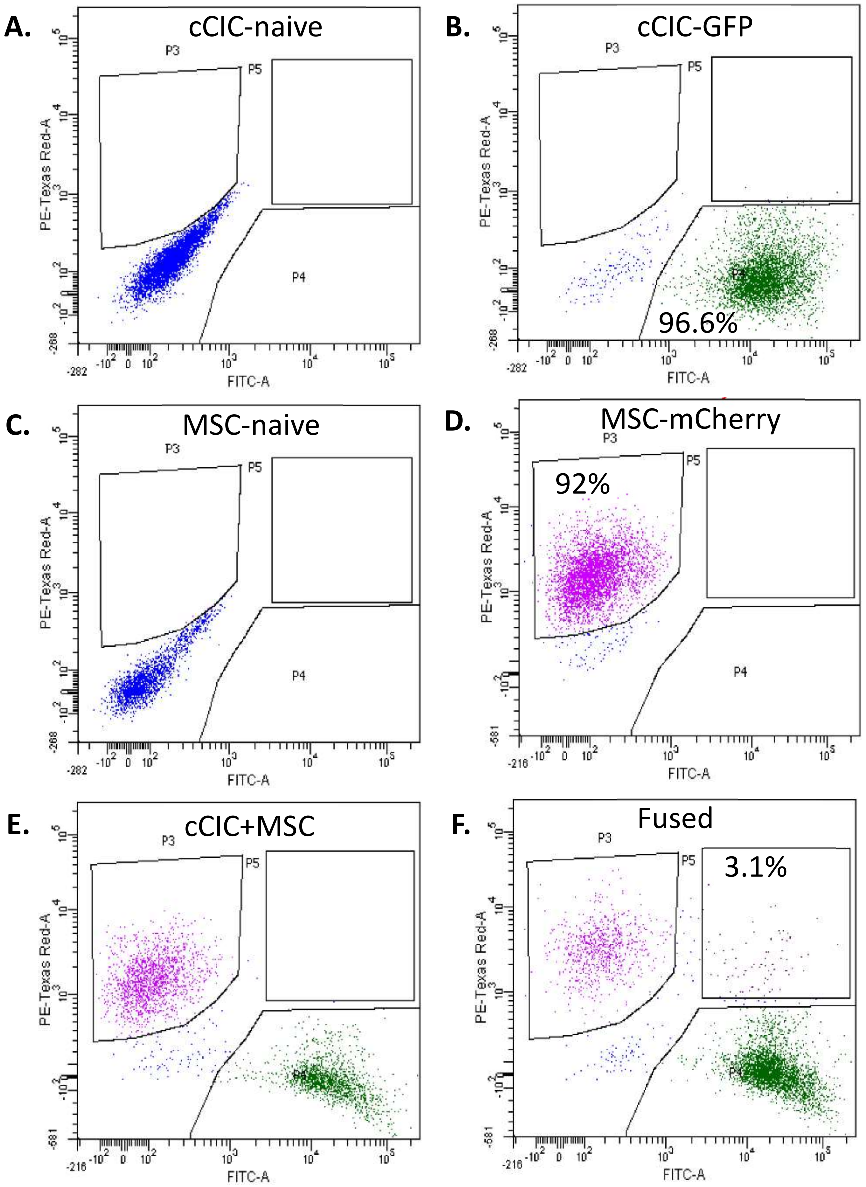


**Supplemental Figure 1:** Flow cytometry plots of one representative fusion experiment including **A.** naive cCICs, **B.** cCIC-GFP, **C.** naive MSCs, **D.** MSC-mCherry, **E.** the combinatorial cCIC and MSC group as the negative control and **F.** Sendai virus-induced fused cells.

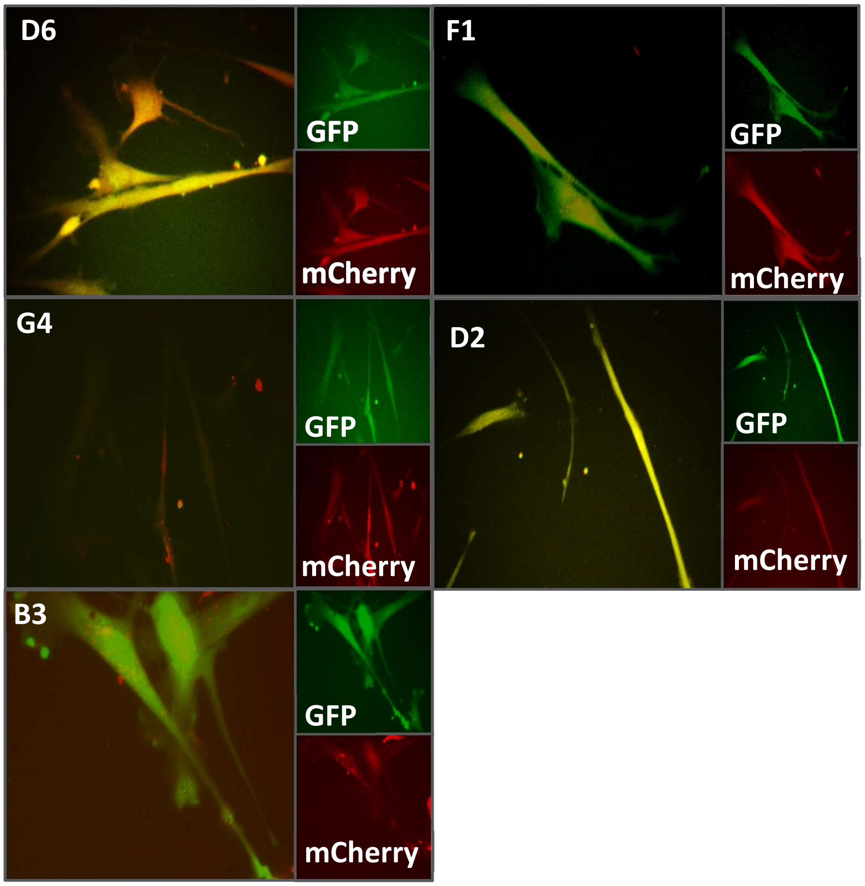


**Supplemental Figure 2:** Live native fluorescent images of hCCs illustrating double positivity of the fused cells.

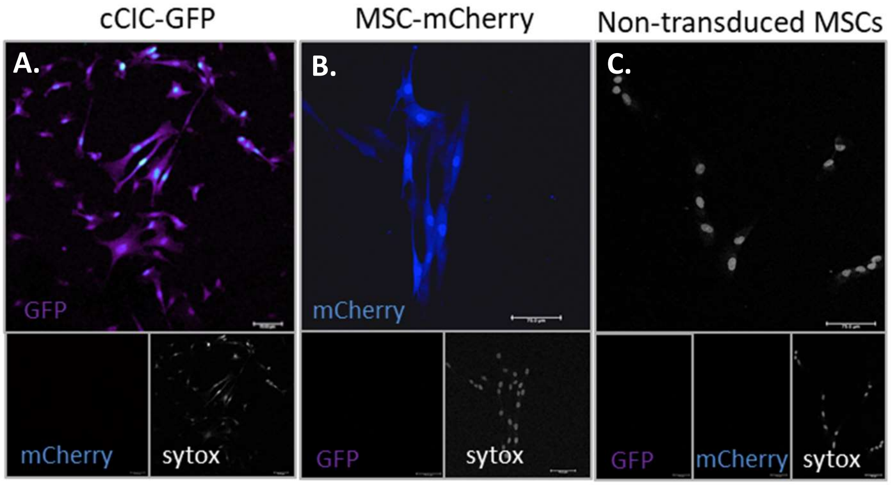


**Supplemental Figure 3:** Immunocytochemistry images of **A.** cCIC-GFP, **B.** MSC-mCherry, and **C.** Non-transduced MSCs stained for GFP and mCherry. Sytox green was used as the nuclear stain.

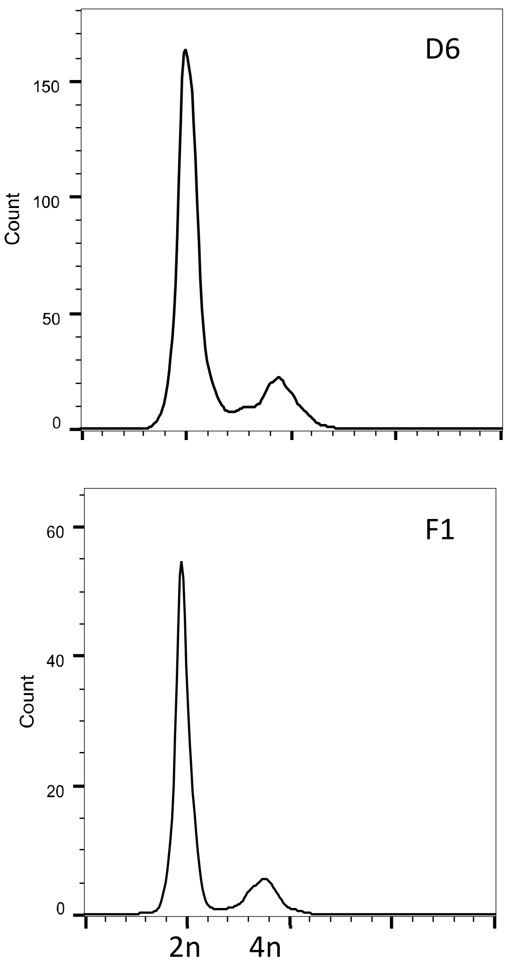


**Supplemental Figure 4:** Flow Cytometry plots for PI/RNAse staining of hCCs D6 and F1.

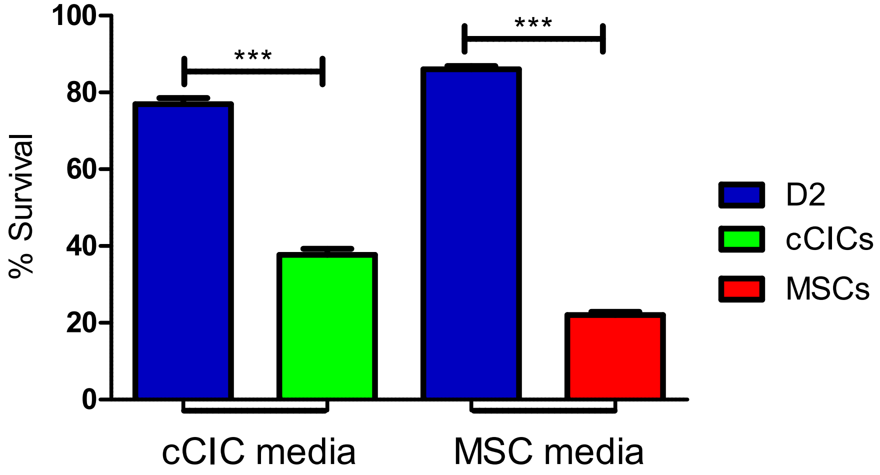


**Supplemental Figure 5:** Percentage of survival (live cells) of D2 clones in cCIC media, cCICs in cCIC media, D2 clones in MSC media and MSCs in MSC media (from left to right) after treatment with hydrogen peroxide (350 µmol/L). Error bars are ± SEM. *** $p<0.001$

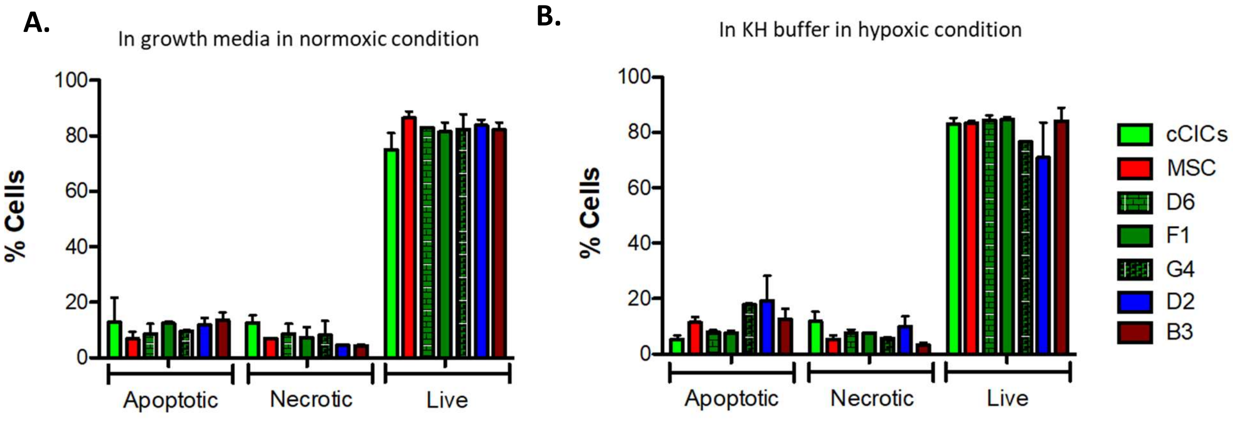


**Supplemental Figure 6:** Percentage of apoptotic, necrotic and live cells **A.** cultured in growth media in normoxic condition and **B.** subjected to KH buffer in hypoxic condition. Error bars are ± SEM.

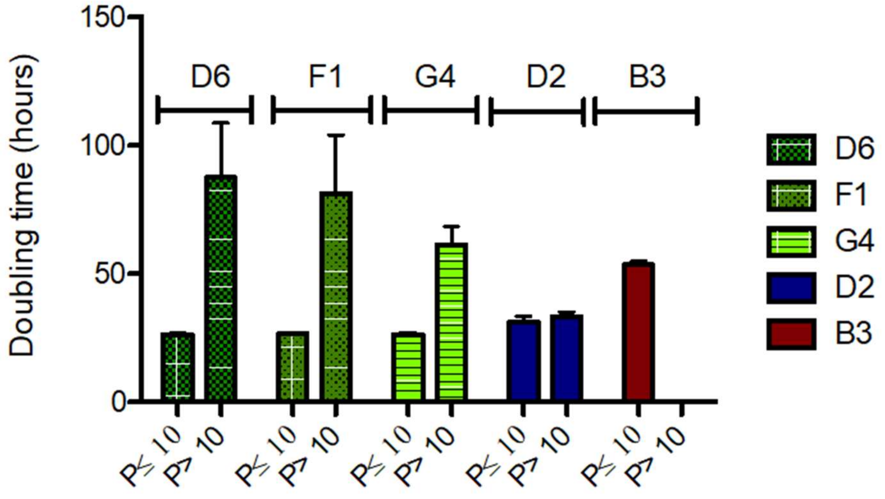


**Supplemental Figure 7:** Cell doubling time of hCCs up to and after passage 10 in culture represented in hours. Error bars are ± SEM.
